# Pleiotropy increases parallel selection signatures during adaptation from standing genetic variation

**DOI:** 10.1101/2024.08.06.606803

**Authors:** Wei-Yun Lai, Sheng-Kai Hsu, Andreas Futschik, Christian Schlötterer

## Abstract

The phenomenon of parallel evolution, whereby similar genomic and phenotypic changes occur across replicated pairs of population or species, is widely studied. Nevertheless, the determining factors of parallel evolution remain poorly understood. Theoretical studies have proposed that pleiotropy, the influence of a single gene on multiple traits, is an important factor. In order to gain a deeper insight into the role of pleiotropy for parallel evolution from standing genetic variation, we characterized the interplay between parallelism, polymorphism and pleiotropy. The present study examined the parallel gene expression evolution in 10 replicated populations of *Drosophila simulans*, which were adapted from standing variation to the same new temperature regime. The data demonstrate that parallel evolution of gene expression from standing genetic variation is positively correlated with the strength of pleiotropic effects. The ancestral variation in gene expression is, however, negatively correlated with parallelism. Given that pleiotropy is also negatively correlated with gene expression variation, we conducted a causal analysis to distinguish cause and correlation and evaluate the role of pleiotropy. The causal analysis indicated that both direct (causative) and indirect (correlational) effects of pleiotropy contribute to parallel evolution. The indirect effect is mediated by historic selective constraint in response to pleiotropy. This results in parallel selection responses due to the reduced standing variation of pleiotropic genes. The direct effect of pleiotropy is likely to reflect a genetic correlation among adaptive traits, which in turn gives rise to synergistic effects and higher parallelism.

## Background

Distinguishing selection from stochastic changes continues to be a major challenge in evolutionary research. The evidence for selection mostly comes from parallel evolution, where similar genes, traits or functions are identified in replicate populations exposed to similar environments [1-4]. Given that non-parallel evolution does not preclude the possibility of selection [5, 6], there has been a sustained interest in elucidating the factors that influence the extent of genetic parallelism (i.e., how similar is the response of the same gene to selection in multiple populations from the same/similar ecological niche(s)) [7].

It has been proposed that the degree of pleiotropy, the number of traits affected by a single gene, may be a potential factor associated with parallelism [7-10]. The majority of hypotheses regarding the influence of pleiotropy on parallel selection responses are (implicitly) related to the ideas of Fisher [11], later modified by Orr [12], focusing on the cost of pleiotropy. This concept posits that any new mutation in a pleiotropic gene results in a shift in trait values away from their optimum across multiple traits. New mutations in highly pleiotropic genes are more likely to have negative fitness effects than mutations in less pleiotropic genes as more traits are displaced from their optima. This “cost of pleiotropy” model [12] is evidenced by the lack of sequence divergence observed in genes with a higher level of pleiotropy [13-16]. Consequently, it can be postulated that less pleiotropic genes are more likely to contribute to adaptation over long evolutionary timescales, given that they have a higher probability of accumulating beneficial genetic variants. This results in a more parallel selection response in different populations/species [17].

Fisher’s model of pleiotropy is simplistic in its assumption of uncorrelated traits and uncorrelated selection. It has been proposed that the influence of pleiotropy on the selection response strongly depends on the genetic correlation of the affected traits and the underlying fitness function which connects phenotypic values with fitness [18-20]. If the fitness effects of the affected traits are positively correlated (synergistic pleiotropy), this may result in stronger net selection [21] and consequently in more parallel selection signatures [22]. This implies that the correlation of fitness effects determines whether pleiotropy increases or decreases parallel selection signatures.

The impact of pleiotropy on parallel evolution has typically been considered in the context of long-term adaptation via de novo mutations. However, many studies of parallel evolution have focused on short-term adaptation [e. g. 23, 24-30], where standing genetic variation plays a more significant role for selection responses than de novo mutations. Nevertheless, it remains unclear whether a similar impact of pleiotropy on evolutionary parallelism is to be expected for standing genetic variation.

Indeed, the empirical evidence regarding the role of pleiotropy for intra-specific adaptive responses is somewhat inconclusive. A gene expression study in European grayling populations adapted to different temperature regimes provided compelling evidence for the constraining effect of pleiotropy [31], which supports the cost of complexity concept. A comparison of gene expression changes in natural *Drosophila* populations along a temperature cline and populations evolved in the laboratory to different temperature regimes demonstrated the importance of correlated fitness effects [21]. A direct test of the influence of pleiotropy on parallel evolution was conducted on 19 independent contrasts of stickleback populations [9]. The authors discovered that pleiotropic genes were more likely to exhibit parallel adaptation signatures [9], thereby supporting the importance of synergistic pleiotropy.

Previous studies on pleiotropy and parallelism did not account for polymorphism (standing genetic variation), which plays a central role for selection responses during short-term adaptation. Although natural variation provides the basis for adaptive responses, it is not well-documented how ancestral variation (polymorphism) determines the level of genetic parallelism during short-term adaptation in replicate populations. Moreover, polymorphism of a gene is also affected by its pleiotropic level while the expectations regarding the influence of pleiotropy on the extent of polymorphism varies. The cost of complexity predicts that pleiotropic genes with uncorrelated fitness effects will be subject to stronger purifying selection, which will result in reduced polymorphism. In the case of multiple selected traits with aligned fitness effect, synergistic pleiotropy may facilitate the occurrence of selective sweeps and thereby reducing variation in pleiotropic genes. Alternatively, it may be possible that once pleiotropic alleles have been established in a population, their multivariate fitness effects lead to a balanced polymorphism, which in turn maintains higher levels of variation. Empirical studies have observed a negative correlation between the level of pleiotropy and sequence/expression polymorphism [32, 33]. This suggests that pleiotropy may preferentially reduce variation, either by purifying selection, synergistic effects, or a combination of both.

In this study we employed gene expression analysis to investigate the interplay between polymorphism and pleiotropy on the degree of parallelism of adaptive responses. We assessed the selective response by measuring gene expression differences between populations from different environments, a widely employed approach [e. g. 34, 35-40]. Specifically, we compared gene expression in the ancestral *D. simulans* population with that of flies that had evolved for 100 generations in a novel hot temperature regime. To infer parallel evolution, the gene expression change was measured in 10 replicate populations. By focusing on the evolutionary response, we circumvent the challenge of connecting the phenotype to an unknown fitness landscape. Furthermore, we take advantage of well-established pleiotropy estimates [41-44] to evaluate the influence of pleiotropy on parallel gene expression evolution. The natural variation in gene expression was determined by examining the differences in expression levels among individual flies from the ancestral *D. simulans* population.

We observed significant interplays between pleiotropy, ancestral variation, and parallel selection responses among genes. The parallelism of gene expression evolution is positively associated with pleiotropy but negatively associated with the ancestral variation in gene expression. Given that pleiotropy was also negatively associated with ancestral variation, it is crucial to differentiate between statistical causality and correlation. Causal analysis revealed that pleiotropy affects parallel evolution both directly and indirectly via ancestral variation. It is likely that the indirect effects operate via the reduction of ancestral variation in pleiotropic genes. The direct effects of pleiotropy on parallelism are probably best explained by synergistic pleiotropy which results in a stronger selection response.

## Methods

### Estimating gene pleiotropy

We approximated the pleiotropy of each gene with two alternative estimators, network connectivity and tissue specificity (τ).

τ indicates how specific the expression of a gene is across different tissues. τ was estimated for each gene using the adult male expression profiles on flyatlas2 [45] as: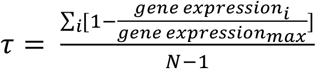, where *N* is the number of tissues examined and *i* indicates each of them *[46]. If* a gene is only expressed in one tissue, τ will equal to 1 while it equals 0 when a gene is expressed at the same level across all tissues. The relationship between τ and pleiotropy is based on the idea that the genes expressed in many tissues are more likely to affect multiple traits than genes expressed in fewer tissues [41, 44, 47, 48]. Hence, we used 1-τ to indicate the pleiotropic effect of a gene.

Network connectivity has also been used as proxy for pleiotropy, as more connected genes are likely to be more pleiotropic [16, 42, 43, 49]. We used published information about the transcriptional regulatory network of *Drosophila*, which was inferred from several sources, including genome-wide chromatin immuno-precipitation, conserved transcription factor binding motifs, gene expression profiles across different development stages, and chromatin modification profiles among several cell types [50]. The connectivity for each gene was estimated as the sum of adjacencies between the focal gene and other genes in the network.

The significant positive correlation (rho=0.54) between 1-τ and network connectivity suggests that both estimates capture a similar, but not identical, signature of the pleiotropic properties among the genes (Supplementary figure 1).

We note that approximating pleiotropic effects in *D. simulans* by using data-rich estimates from the close relative *Drosophila melanogaster* could potentially compromise the interpretability of our results. However, comparative analyses of tissue-specific gene expression have shown an extremely high consistency across species in animals [51] and plants [52, 53]. Hence, these data suggest that at least for tissue specificity, one of the pleiotropy measures used, estimates from one species can be safely used for close relatives.

### Experimental evolution and common garden experiment

The procedures of the evolution and common garden experiments were described in [5, 35, 54]. Briefly, 202 isofemale lines collected from Florida, USA were used to constitute 10 outbred *Drosophila simulans* populations, which have been exposed for more than 100 generations to a laboratory experiment at 28/18 °C with 12hr light/12hr dark photoperiod. The census population size of each replicate population is 1000 to 1250 adult flies.

Each founder isofemale line was maintained at a small population size (typically less than 50 individuals) at 18 °C on standard laboratory food. Adaptation to the laboratory environment with the residual heterogeneity or de novo mutation is unlikely due to the small effective population size during the maintenance of each line [5]. This is supported by the observation that 50 generations of maintenance as isofemale lines does not result in significant genetic differentiation from the ancestral population [55]. Furthermore, mutations that occur in the laboratory are mostly recessive [56], and the effects are likely to be masked because after two generations of random mating during common garden maintenance, most individuals will be heterozygous for isofemale line-specific variants.

The collection of samples for RNA-Seq was preceded by two generations of common garden experiment (CGE). As described previously [5, 35, 54], at generation 103 of the evolution experiment, five biological replicates of reconstituted ancestral populations were generated by pooling five mated females from 184 founder isofemale lines [57] and reared together with 10 independently evolved populations for two generations with controlled egg density (400 eggs/bottle) at the same temperature regime as in the evolution experiment. Three biological replicates were included for each independently evolved population (each biological replicate was cultivated separately for the two generations of common garden). From each biological replicate, a sample of 50 five-day-old adult males were collected (Figure 1a).

**Figure 1.**
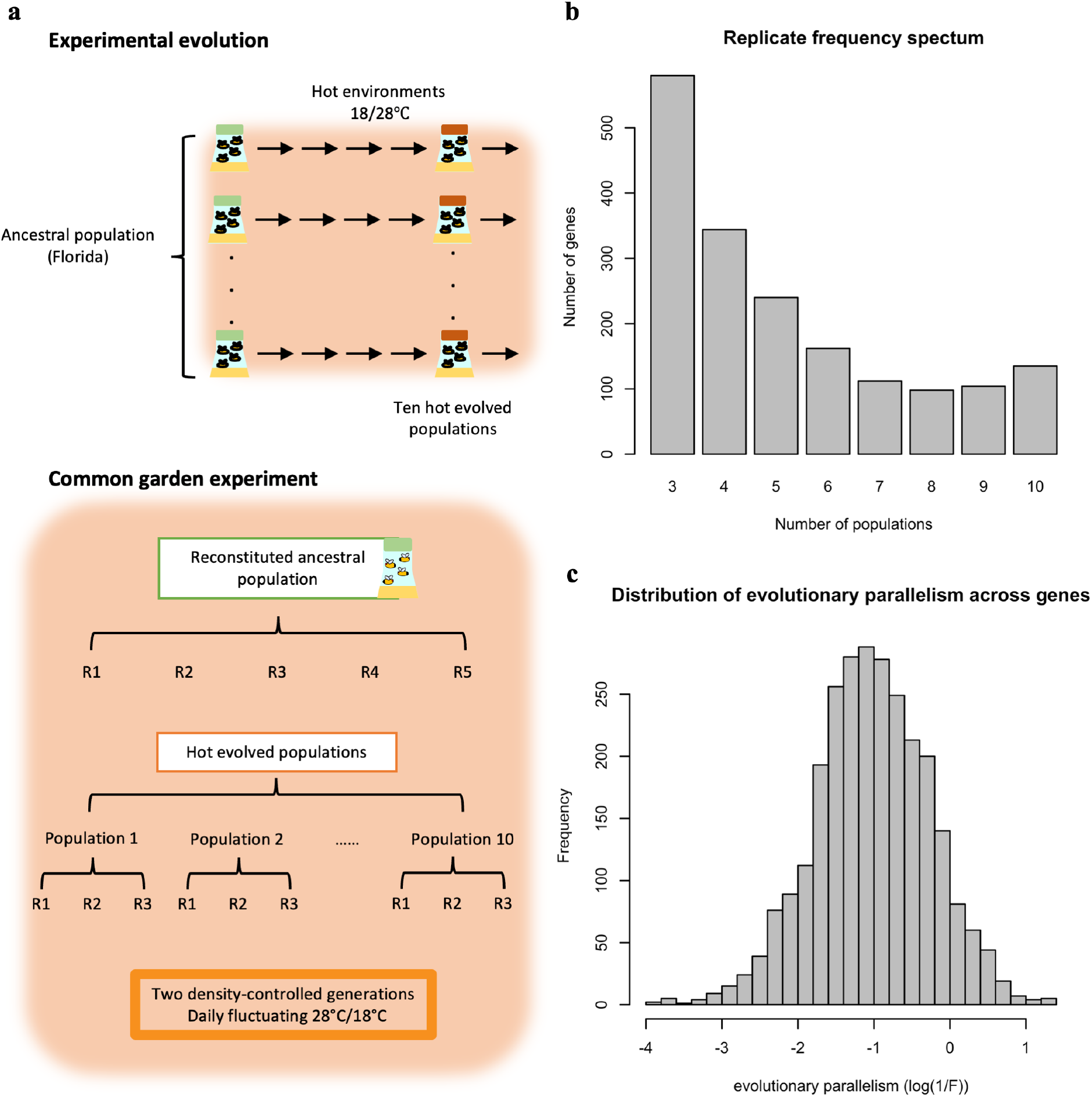
Overview of the experimental procedures (a) and the variation in parallel gene expression evolution across genes (b,c). **a.** Experimental Evolution: ten replicated populations seeded from one common founder population have been evolving for >100 generations in a hot laboratory environment at 18 and 28°C. Common Garden Experiment: at the 103^rd^ generation of the evolution experiment, the ten evolved populations (each with three biological replicates) were maintained together with the reconstituted ancestral population (with five biological replicates) for two generations in the same environment as in the evolution experiment. After two generations in common garden, 50 males from each biological replicates were pooled and subjected to RNA-Seq. **b**. Replicate frequency spectrum. Number of populations (x-axis) in which a given gene experienced a significant change in gene expression. The y-axis indicates the number of genes in each category. Most of the genes experienced a significant change in gene expression in few evolved populations while much fewer genes were significant in all 10 populations. This pattern suggests that the parallelism in gene expression evolution differs across genes. **c**. The distribution of gene expression evolution parallelism (log(1/F)) across genes. Larger log(1/F) values indicate more parallel evolution of a gene. The exhibit variation suggests that genes varied in their parallelism of its expression evolution.

### Quantifying the evolutionary parallelism in gene expression across ten replicate populations

The raw RNA-Seq count table of five replicates of the reconstituted ancestral population and 10 independently evolved populations, each with three biological samples, was taken from (Jaksic et al., 2020). The raw read counts of each gene were normalized with the TMM method implemented in edgeR [58]. Only genes with at least 0.1 normalized counts per million base (CPM) across all samples were considered for further analysis. Because we are interested in the evolutionary response in each independently evolved population, we contrasted the three biological samples from each evolved population to the five biological samples from the ancestral population. The differential expression (DE) analysis was done separately in each of the ten evolved populations. For DE analysis, we utilized negative binomial generalized linear modeling implemented in edgeR to fit the expression to the model *Y = E + ε*, in which *Y* stands for gene expression, *E* is the effect of evolution and *ε* is the random error. Likelihood ratio tests were performed to test the effect of evolution. P-values was adjusted using Benjamini-Hochberg false discovery rate (FDR) correction [59]. After we identified DE genes in each evolved population, we constructed a replicate-frequency spectrum (RFS; [5]). We considered genes with significant changes in the same direction in at least three evolved populations as putatively adaptive genes.

To quantify the degree of parallel gene expression evolution, we calculated the expression changes in response to evolution (log_2_FC; fold change in expression between ancestral and evolved sample; see below) of each putatively adaptive gene (n=1,775) in each evolved sample.

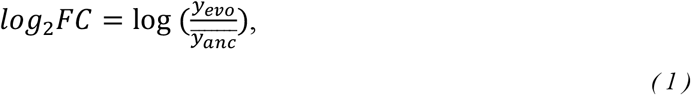

where *y*_*evo*_, is the expression value (CPM) per gene per evolved sample, and 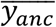 is the overall mean expression value (CPM) per gene across five ancestral samples.

We calculated the ratio of between-evolution population variation (MS_pop_) and residual (MS_e_) among the expression changes (log_2_FC), which is denoted as F.

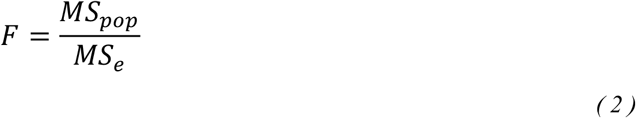

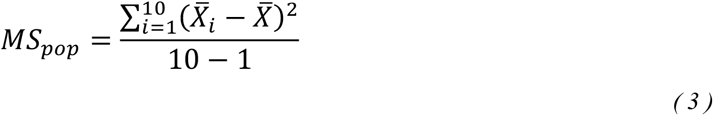

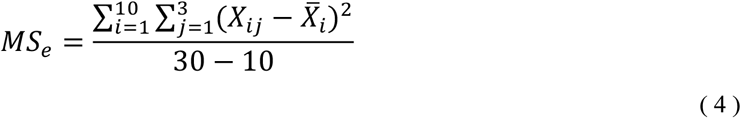

where *X*_*ij*_ *represen*t the log_2_FC value for the j^th^ biological sample in the i^th^ evolution population. i=1, 2, …,10, j=1, 2, 3.

As F reflects the heterogeneity of the evolutionary response for a gene across ten evolutionary populations, we used the reciprocal of F (i.e., 1/F) to quantify the degree of parallelism in gene expression evolution.

### Estimating gene expression variation in an outbred ancestral population

The variation in expression of each gene within an outbred ancestral population was estimated from 20 adult males sampled from an outbred ancestral population [60]. We distinguished biological variation and measurement error of each gene using the statistical method implemented in edgeR [58], where the biological variance across individual (biological coefficient of variation; BCV; defined in [58]) of each gene can be estimated (tag-wised dispersion). BCV^2^ was used to represent the expression variance relative to the mean of each gene within the population. To illustrate that the gene expression variation is not explained by random measurement error, we took the data from two ancestral population replicates, which are considered to be genetically identical and calculated BCV^2^ of expression levels across individuals for each population replicates separately. We found that both mean and variance in expression for each gene was similar in two ancestral population replicates (mean: rho>0.99; variance: rho=0.8, p-value<2.2e-16), suggesting that sequencing noise does not mask the biological variability.

### Causal analysis

To infer the causal relationship between three factors: pleiotropy (Pl), ancestral variation (A) and parallelism (Pa), we applied causal analysis. The causal analysis is built upon a previously published statistical framework [61]. We consider five possible causal relationships between pleiotropic effects, ancestral variation and parallelism (Figure 5). Our goal is to determine which of the five models is best supported by the data. The selection is based on Bayesian information criteria (BIC) which are calculated from maximum likelihood of the respective models. The likelihood of each model is represented as:

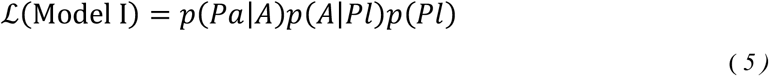

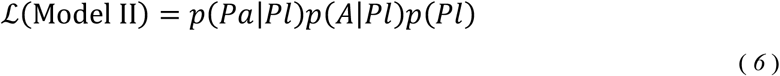

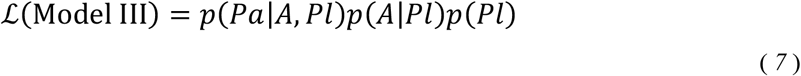

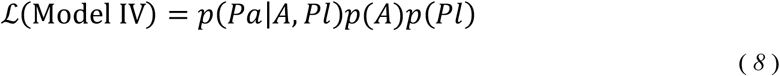

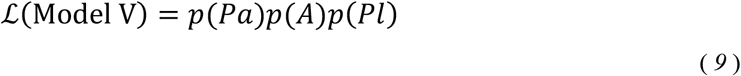

p(Pl) is the probability distribution of the pleiotropic effect. To utilize normal probability density function in the derivation of the likelihoods for each joint probability distributions, we transformed the random variables Pa and A to be normally distributed (natural log transformation on 1/F and BCV^2^). The exact forms of these likelihoods are given in supplementary information.

For each model, we maximized the likelihood and estimated the parameters using standard maximum likelihood methods. Based on the maximum likelihood, we computed the BICs for each model; the model with the smallest BIC is the one best support by the data.

BIC is calculated as:

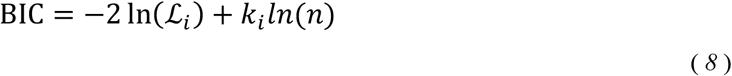

Where *ℒ*_*i*_ is the likelihood for model I-V, *k*_*j*_ is the corresponding number of free parameters and n is total number of putatively selected genes. The analysis was performed independently on two measures of pleiotropy (tissue specificity and network connectivity).

### Estimating the sizes of direct and indirect pleiotropic effects on parallelism

We used path analysis to estimate the effect sizes of direct and indirect pleiotropic effects on parallelism [62]. We standardized all three transformed variables (Parallelism (Pa), Pleiotropy (Pl), Ancestral variation (A); see supplementary information) by subtracting the mean and dividing by the standard deviation for the following analysis.

First, we fit the regression model for all three variables across all putatively adaptive genes:

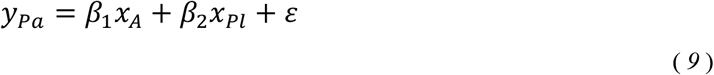

where *y*_*pa*_ stands for parallelism, *x*_*A*_ is ancestral variation, *x*_*pl*_ is pleiotropy measure and *ε* is random error. *β*_*1*_ and *β*_*2*_ are the regression coefficients corresponding to *x*_*A*_ and *x*_*pl*_, respectively.

Second, we fit another regression model for ancestral variation and pleiotropy measure across all putatively adaptive genes:

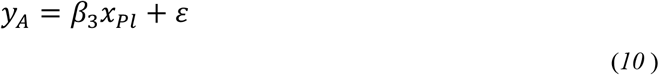

where *y*_*A*_ stands for ancestral variation, *x*_*pl*_ is pleiotropy measure and *ε* is random error. *β*_*3*_ is the regression coefficient corresponding to *x*_*pl*_.

The sizes of the direct effect of pleiotropy on parallelism will be *β*_*2*_. The size of the indirect effect of pleiotropy on parallelism via ancestral variation will then be the product of *β*_*1*_ × *β*_*3*_.

The analysis was performed independently on two different measures of pleiotropy.

### Computer simulations

In our interpretation of the indirect effect of pleiotropy on parallelism, we assumed that ancestral variation in gene expression is negatively correlated with its parallel evolution. Because this has not been previously demonstrated, we used computer simulations to illustrate how the level of standing genetic variation impacts the parallelism of adaptive responses after a shift in trait optimum. For simplicity, we considered four redundant genes equally contributing to a fitness-associated trait. The phenotype (expression level) of each gene is controlled by a different number of genetic loci (5, 15, 30 and 50) with equal effect. The underlying assumption is that these four genes are redundant with in terms of their fitness effect during the experimental evolution, but they exhibit different levels of pleiotropy and thus have historically experienced different levels of purifying selection, leaving different levels of genetic variation at the start of the experiment. Using mimicrEE2 [63], we evolved 10 replicate populations (N=300) derived from the same set of 189 natural *Drosophila* haplotypes [5] in same selection regime with a shift in trait optimum of one standard deviation relative to the ancestral phenotypic distribution. After 100 generations, we determined the evolutionary parallelism of the four genes using the same approach as for the empirical data. Each simulation was repeated 100 times and one out of four genes was randomly picked from each simulation run to generate Figure 4.

## Results

### The parallelism of expression evolution differs among genes

We investigated the degree of parallelism in gene expression evolution, using an RNA-Seq dataset from an experimental evolution study in which 10 replicated populations with the same genetic variation were exposed to the same environment for more than 100 generations [29, 54, 60] (Figure 1a). While previous studies have focused on the shared parallel response across replicated populations [29, 54, 60], we re-analyzed the RNA-Seq data to quantify the degree of evolutionary parallelism across different genes. We identified 1,775 putatively selected genes with a significant change in gene expression in the same direction in response to the novel environment in at least three populations. This conditioning on multiple replicate populations with a significant change in the same direction reduces false positives due to drift [9, 64]. Relaxing the threshold of responding populations provides qualitatively similar results.

Barghi et al. (2019) introduced the replicate frequency spectrum to describe the parallel genomic selection response in replicate populations. The replicate frequency spectrum of DE genes shows considerable heterogeneity among populations, with many genes displaying significant gene expression changes only in a few populations (Figure 1b). The replicate frequency spectrum is strongly influenced by the significance threshold used and provides an incomplete quantitative measure of parallelism. To obtain a more quantitative measure of parallelism, we evaluated the between-population variation in the evolutionary response (log_2_FC) of each putatively adaptive gene (n=1,775) while controlling for measurement error (F). The parallelism of evolution for each gene is then quantified as 1/F (Figure 1b; see Materials and Methods). A higher value of 1/F indicates a more parallel evolutionary response among replicated populations (Supplementary figure 2). Importantly, the measure of parallelism varies between genes (Figure 1c) and is positively correlated with the number of populations in which a significant change in gene expression is observed for each gene (Supplementary figure 3).

### Pleiotropy is positively associated with the degree of parallelism in gene expression evolution

Pleiotropy is one of the factors that influence the degree of parallel evolution [7-10]. Therefore, we tested whether and how the pleiotropic effect of a gene influences the degree of parallelism of its expression evolution in a new environment. We correlated two different measures of pleiotropy (1-τ(tissue specificity) and network connectivity; see materials and methods) of each putatively adaptive gene (n=1,775) with the parallelism of its evolutionary responses (1/F). We observed a significant positive correlation between the parallelism and the degree of pleiotropy, (rho=0.26, p-value<2.2e-16 for 1-τ and rho=0.11, p-value<5.6e-07 for network connectivity) (Figure 2 and Supplementary figure 4a). This suggests that pleiotropy may enhance parallel changes in gene expression during evolution.

**Figure 2.**
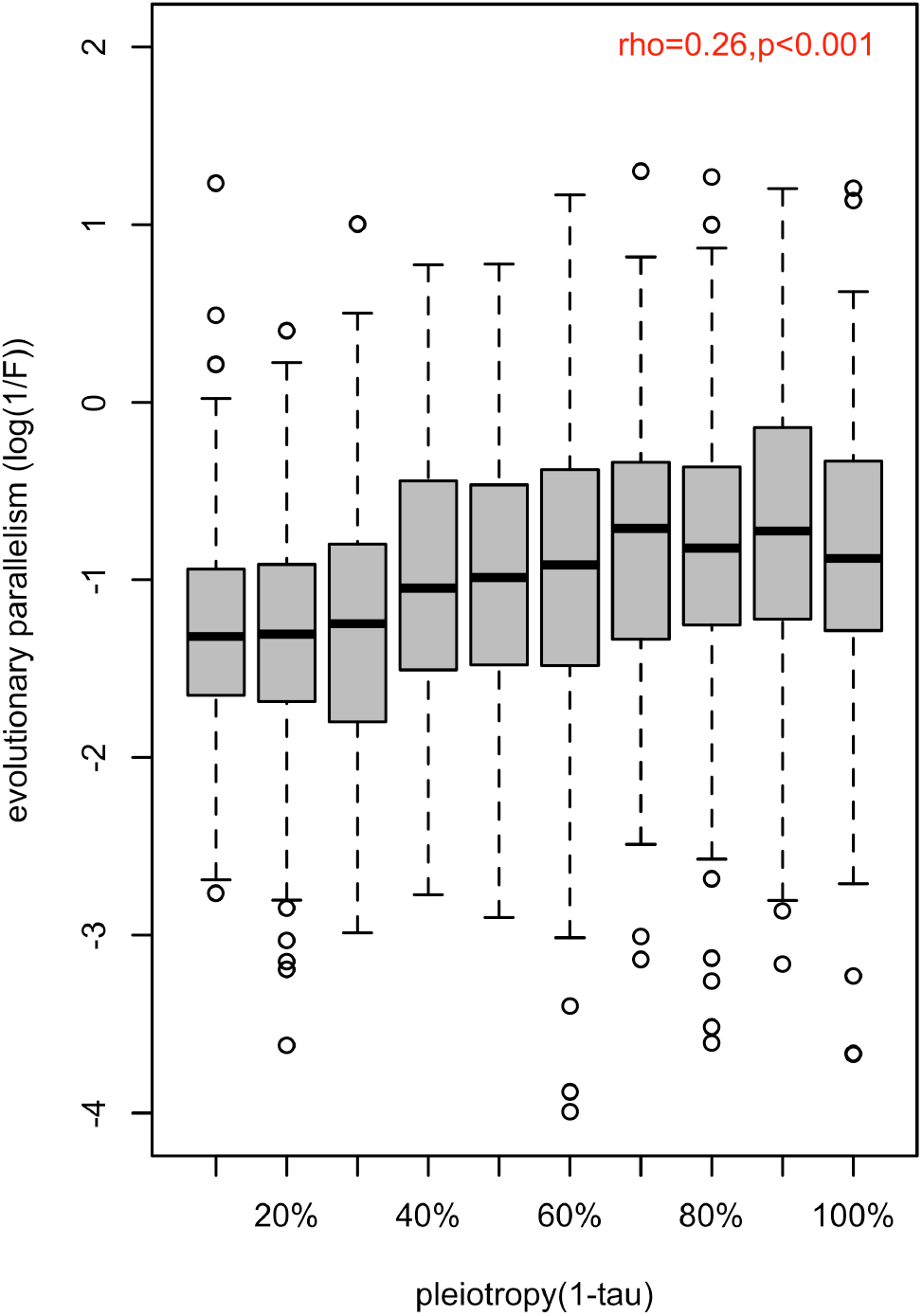
Association between the magnitude of evolution parallelism and the strength of pleiotropy. The distribution of evolution parallelism (log(1/F)) of different genes is shown in boxplots binned by their strength of pleiotropy (1-*τ*). Overall, the strength of pleiotropy was positively correlated with evolution parallelism (*ρ*=0.26, p-value<2.2e-16).

### Pleiotropy constrains the ancestral (natural) variation in gene expression

The negative effect of pleiotropy on genetic variation over longer time scales, mostly between species, is well documented [13-16], but it not yet clear whether the same effect is seen for adaptation from standing genetic variation. We reconstituted the ancestral population from 189 isofemale lines [60] and estimated the expression variance (BCV^2^; see materials and methods) of each putatively adaptive gene (n=1,775) in two replicate sets of 20 individuals. Since the variance in gene expression is positively correlated with genetic variation in cis-regulatory regions [65] and the heritability of gene expression is high [66], we assumed that the observed variation in gene expression has a genetic basis, rather than being noise (see discussion). We observed a highly significant negative correlation between the level of pleiotropy and the magnitude of ancestral variation in gene expression (Figure 3a and Supplementary figure 4b; rho=-0.34, p-value<2.2e-16 for 1-*τ* and rho=-0.37, p-value<2.2e-16 for network connectivity). To exclude that this correlation was driven by our filtering for significant expression changes in at least three populations, we repeated these analyses with all genes (n=9,882) and also observed a significant correlation (rho=-0.45, p-value<2.2e-16 for 1-*τ* and rho=-0.38, p-value<2.2e-16 for network connectivity, data not shown). Hence, we suggest that the negative relationship between pleiotropy and polymorphism/diversity [32, 33] (i.e. reduced variants, lower frequencies and/or lower allelic effects) can be extended to gene expression.

**Figure 3.**
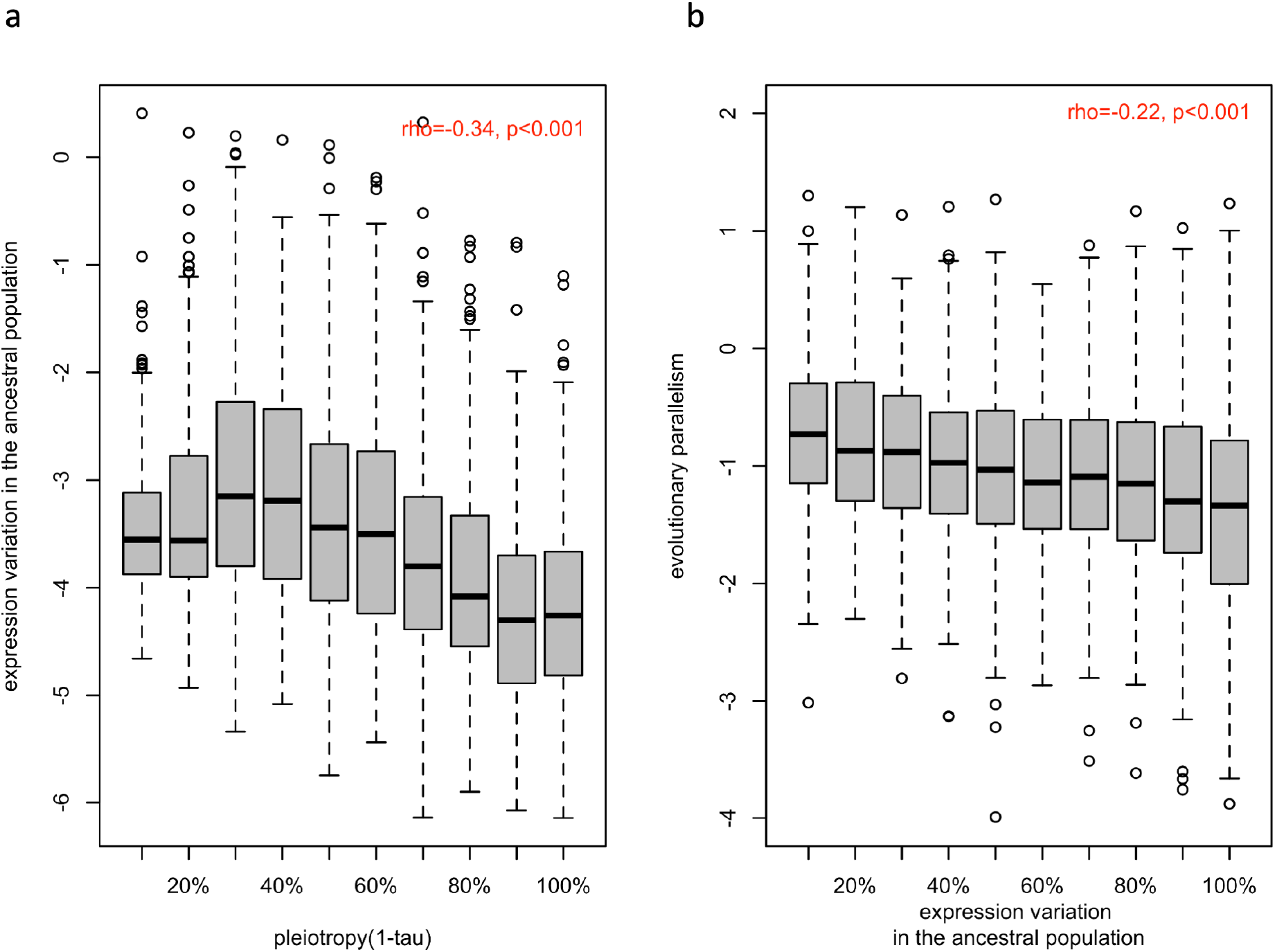
Association between ancestral variation in gene expression with the strength of pleiotropy (1-*τ*) (a) and with the evolution parallelism (b). **a**. The distribution of gene expression variation in the ancestral population (log (BCV^2^)) of different genes is shown in boxplots binned by their strength of pleiotropy (1-*τ). Overa*ll, the strength of pleiotropy was negatively correlated with ancestral variation in gene expression (*ρ*=-0.34, p-value<2.2e-16). **b**. The distribution of evolutionary parallelism (log(1/F)) of different genes is shown in boxplots binned by their strength of ancestral variation. The strength of parallelism is negatively correlated with the ancestral variation in gene expression (ρ=-0.22, p-value<2.2e-16).

**Figure 4.**
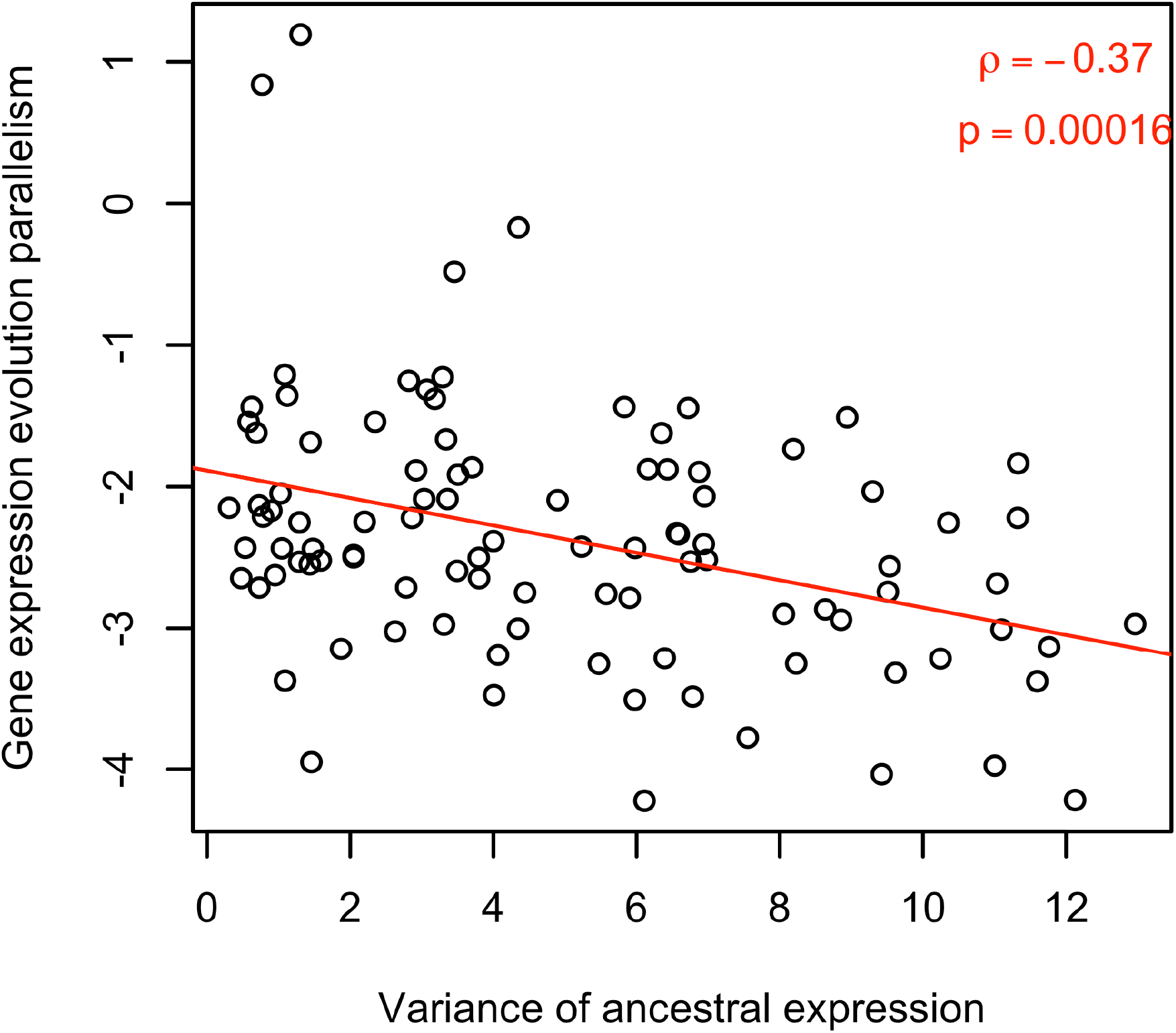
Computer simulations illustrate the negative effect of ancestral variation on parallel evolution. Computer simulations assume a fitness related trait determined by the expression of four genes. The expression of these four genes is determined by a different number of genetic loci, reflecting the influence of pleiotropy on ancestral genetic variation. A shift in trait optimum was used to illustrate the connection between ancestral variation in expression (x-axis) and parallelism of expression change across ten evolved populations (y-axis). The negative correlation between them (rho = -0.26, p-value < 1.77e-07) suggests that the gene with less ancestral variation are resulting in more parallel responses.

### Evolutionary parallelism is negatively associated with ancestral variation

For a better understanding of the factors determining parallel evolution, we examined the influence of ancestral variation in gene expression on the parallel evolution of gene expression. We observed a significant negative correlation between ancestral variation and parallelism of the evolutionary response (rho=-0.22, p-value<2.2e-16; Figure 3b). We suggest that this pattern could be explained by genetic redundancy [5, 67], which describes the phenomenon that a polygenic trait can shift its phenotypic value through different combinations of contributing loci: When the diversity of the contributing loci is reduced (either through fewer loci, lower ancestral frequencies or smaller variation in effect size), fewer combinations of the contributing loci are available for adaptation, resulting in more parallel adaptive responses. To confirm this, we performed computer simulations of replicated polygenic adaptation and found that lower ancestral variation (i.e. fewer contributing loci) leads to more parallel trait evolution (Figure 4). Hence, that natural variation in the ancestral population could be one of factor determining the parallel adaptive response in replicate populations. Although we focused on the phenotypic response to match gene expression, it is interesting to note that a similar result was obtained for parallelism at the genetic level with different levels of standing genetic variation [68]. If pleiotropy reduces variation, the positive correlation between pleiotropy and parallelism may be an indirect (correlative) effect, rather than a direct consequence of pleiotropy.

### Testing causality

Our previous analyses indicated that parallelism is affected by pleiotropy and ancestral variation. Because pleiotropy is negatively correlated with ancestral variation for both pleiotropy measures (Figure 3a and supplementary figure 4b), it is not clear whether pleiotropy affects parallelism directly or indirectly through the negative correlation with ancestral variation. We used causal analysis, which based on a Bayesian statistical framework [61], to disentangle the causal relationship between the three different parameters. Since the directionality of pleiotropic effects on ancestral variation has been previously demonstrated, and ancestral variation is unlikely to influence the pleiotropy of a gene, we considered five possible causal relationships between pleiotropic effects, ancestral variation, and parallelism (Figure 5). In the first model, parallelism is determined by ancestral variation, which is shaped by pleiotropic, but pleiotropy has no direct effect on parallelism. In model II, the pleiotropic effect affects ancestral variation and parallelism independently, but ancestral variation has no causal effect on parallelism. In model III, pleiotropy affects both ancestral variation and parallelism, and ancestral variation also determines parallelism. In model IV, parallelism is determined by ancestral variation and pleiotropy, but pleiotropic effects are assumed to have no effect on the ancestral variation. This serves as a null model to confirm the indirect effect of pleiotropy on parallelism via ancestral variation. We also add a global null model with no correlation between these three factors (model V, Figure 5). The analysis was performed independently for two measures of pleiotropy (tissue specificity and network connectivity).

**Figure 5.**
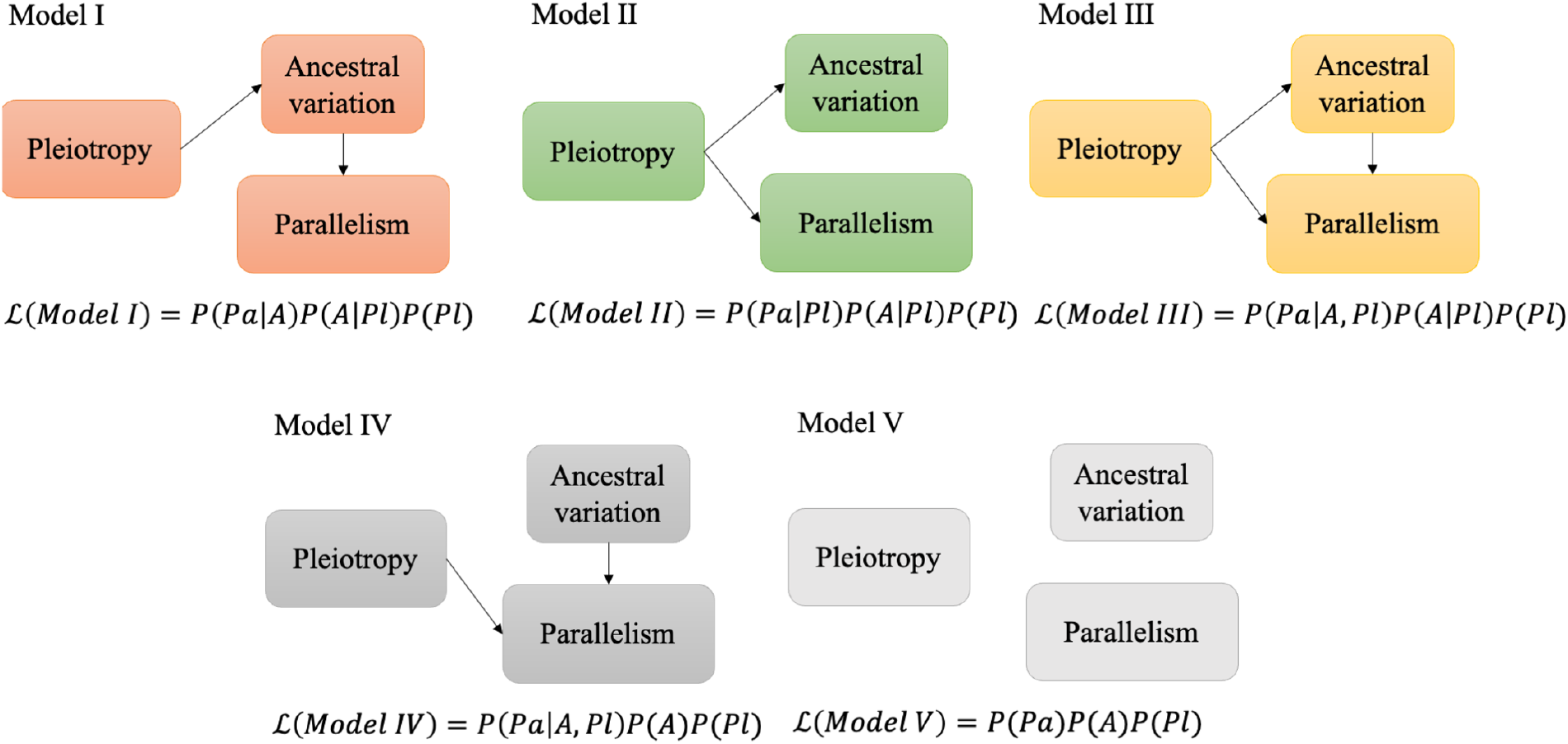
Schematic figure illustrating the causal models evaluated. Five possible relationships between pleiotropic effects, ancestral variation, and evolutionary parallelism. *ℒ* denotes the likelihood of each model given the data. P is the probability or conditional probability of the measurements for each gene; *Pa* is the parallelism level and *A* is the level of ancestral variation in gene expression. *Pl* is the pleiotropic effects. See materials and methods for a more detailed description.

For tissue specificity, our data were better explained by model III (i.e., model III has the lowest BIC, Table 1), suggesting that both direct and indirect effects of pleiotropy determine the degree of parallelism of adaptive gene expression evolution. On the other hand, when pleiotropy was estimated by network connectivity, model I had a slightly lower BIC than model III (Table 1). Thus, our causal analysis does not provide evidence for a direct effect of network connectivity. We further quantified the strength of the direct and indirect effects of pleiotropy on parallelism using a path analysis (see materials and methods). The path analysis confirmed the significant direct effect of tissue specificity, but network connectivity has no statistically significant direct effect on the parallelism of gene expression evolution (Table 2 and Table S1). Although the relevance of direct effects of pleiotropy on parallel gene expression evolution differs for the two pleiotropy estimates, the similarity of the indirect effect sizes for both pleiotropy measures is striking (Table 2). Taken together, these results suggest that pleiotropy may enhance the parallel evolutionary response of gene expression directly and indirectly through its influence on ancestral variation.

**Table 1.**
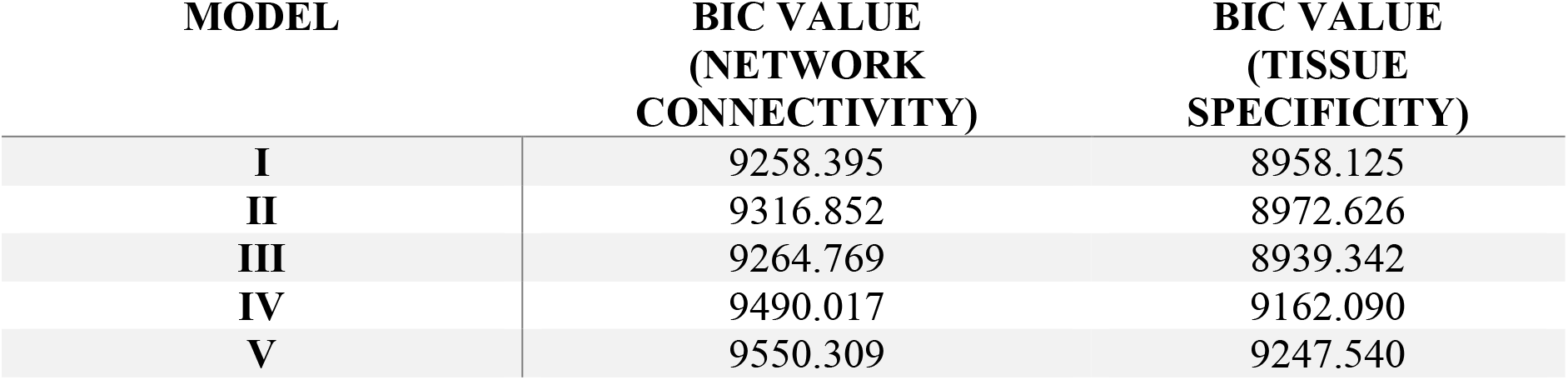
BIC value for Model I-V. The model with the smallest BIC is the one best support by the data.

| **MODEL** | **BIC VALUE**<br>**(NETWORK**<br>**CONNECTIVITY)** | **BIC VALUE**<br>**(TISSUE**<br>**SPECIFICITY)** |
| --- | --- | --- |
| **I** | 9258.395 | 8958.125 |
| **II** | 9316.852 | 8972.626 |
| **III** | 9264.769 | 8939.342 |
| **IV** | 9490.017 | 9162.090 |
| **V** | 9550.309 | 9247.540 |

**Table 2.**
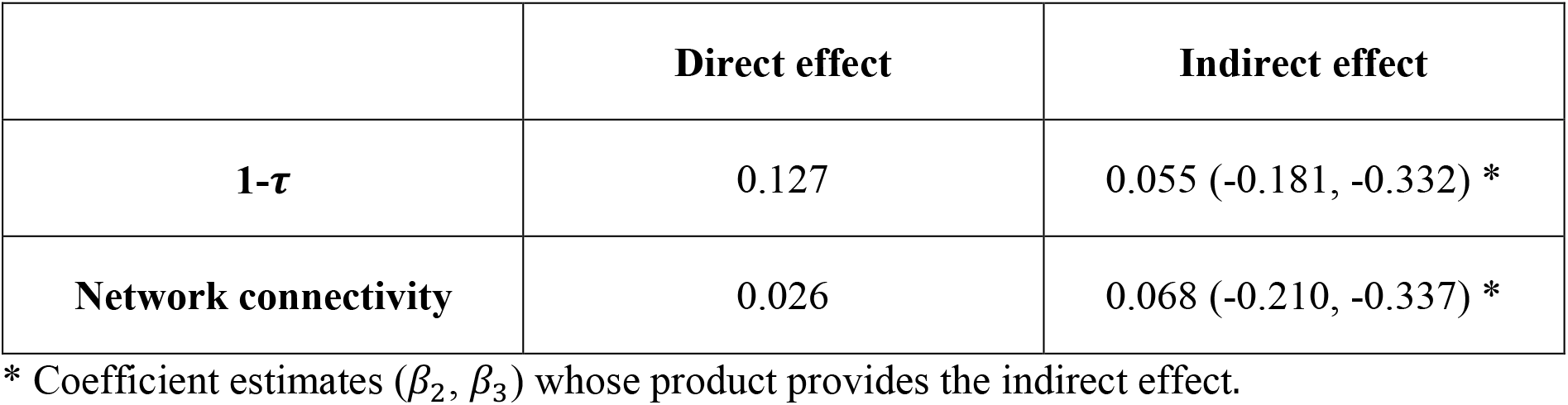
The size of direct and indirect pleiotropic effects on the evolutionary parallelism.

## Discussion

This study examined the influence of pleiotropy on the parallelism of the adaptive response from standing genetic variation. By examining the role of natural variation, which is important for parallelism of short-term evolution, we extended the scope of previous studies. Because differences in genetic background and selection strength among replicated populations cannot be controlled in natural populations and may affect the degree of parallelism, we relied on experimental evolution for our study. We took advantage of ten replicated populations that have evolved from the same genetic background in the same hot environment for more than 100 generations [5].

The adaptive response during adaptation to the new hot temperature regime was measured by changes in gene expression. This approach has the advantage that pleiotropic effects and their associated fitness function, which are typically not known, are already included in the adaptive response. Furthermore, we were able to rely on two well-established measures of pleiotropy to study the interplay with parallelism and natural variation.

A key finding of our study is that that adaptation from standing genetic variation results in complex interplays between pleiotropy, ancestral variation, and parallelism. Of particular interest is the observed negative correlation between pleiotropy and ancestral variation in gene expression, which confirms previous observations [32, 33]. This suggests that the primary effect of pleiotropy is not to maintain variation. Rather, stronger purifying selection removes pleiotropic variants from the population, resulting in a reduced standing genetic variation. The reduced variation in more pleiotropic genes could reflect historical hard sweeps or purifying selection. A similar pattern is seen over longer evolutionary timescales: less sequence divergence in pleiotropic genes [13-16].

An important difference between adaptation from de novo mutations and standing genetic variation concerns the implications for parallel evolution. While pleiotropic genes are less likely to contribute to parallel evolution from de novo mutations [7], our results show that adaptation from standing genetic variation favors parallel selection responses of pleiotropic genes (Figure 2, Supplementary Figure 4a). It is important to note that many previous studies focused on sequence variation, whereas we used expression variation. While it is possible that these two types of variation behave differently, we do not believe that this is the case as sequence polymorphism and expression variation are correlated [69]. Furthermore, a recent study focusing on genomic changes [9] reached similar conclusions.

A very important assumption in our interpretation is that differences in gene expression variation in the ancestral population reflect regulatory sequence polymorphism rather than noise. The high correlation in the variance of gene expression between two replicate groups of individuals indicates that our measures do not reflect technical noise but have a biological basis. However, stochastic fluctuations in gene expression have also been observed among cells in single-celled and multicellular organisms and the variance in gene expression differs among genes [70-72]. In this case, the differences in gene expression variance are biological in nature, but not associated with sequence variation. We do not believe that variation in the control of gene expression levels among genes can explain our pattern because *Drosophila* is a multicellular organism and we used whole body RNA-Seq data for our analysis. Therefore, even if expression is controlled to different extents among genes, by averaging across multiple cells, this does not affect our results. The only scenario we can imagine in which gene expression noise could propagate across the many cells of a multicellular organism is that during early development, differences in gene expression of a key regulator could affect downstream gene expression throughout the entire organism. However, we do not think this is a likely scenario, especially in the light of previous results showing that genetic variation in cis-regulatory regions correlates with expression variation [65]. Furthermore, whole-body gene expression in *Drosophila* is highly heritable, suggesting that gene expression is not stochastic [66]. Nevertheless, we would like to point out that our observations are valid irrespective of whether the correlation between parallelism and expression variation is the result of genetic polymorphism or genetic control of expression noise.

Based on the significant correlation results, the question arose as to whether the positive correlation between pleiotropy and parallelism, is causal (direct effect) or merely correlational (indirect effect). Regardless of the measure of pleiotropy, our causal analysis showed a consistent indirect effect of pleiotropy on parallelism (Table 1, 2). The estimated indirect effects were very similar for both measures of pleiotropy (Table 2). This indirect effect of pleiotropy could be explained by the reduced variation in the expression of more pleiotropic genes in the ancestral population leading to a more parallel selection response of gene expression. For the direct effect, however, the results differed between the two measures of pleiotropy. Only tissue specificity had a significant direct effect, which was even larger than the indirect effect (Table 2). No significant direct effect was found for network connectivitiy. The discrepancy between the two measures of pleiotropy is particularly interesting given their significant correlation (Supplementary Figure 1). This suggests that both measures capture aspects of pleiotropy that differ in their biological implications. The direct effects of pleiotropy, most likely arise from the positively correlated fitness effects of non-focal traits (synergistic pleiotropy). The synergistic pleiotropic effects result in a stronger fitness advantage, which translates into a more parallel selection response - as previously shown [22]. Given that temperature is one of the most important abiotic factors, it is expected that positive genetic correlations between temperature-adaptive traits have been selected over long evolutionary timescales, probably prior to the speciation of *D. simulans*. Thus, it is conceivable that positive pleiotropy for temperature adaptation related traits is common and thus determined the positive contribution of pleiotropy to the parallel evolution observed here.

We focused our analysis on genes that exhibited evidence of adaptive responses, defined as significant changes in the same direction across at least three populations. This approach allowed us to investigate the extent of parallel adaptive responses and how pleiotropy contributes to variation in parallelism among genes. Interestingly, when comparing pleiotropy between DE genes (those showing adaptive responses) and non-DE genes (those that did not change their response during evolution), we observed significantly higher pleiotropy in the non-DE gene group (Supplementary figure 5). This finding suggests that extremely high levels of pleiotropy may constrain evolutionary responses. Overall, the pattern of increased pleiotropy associated with evolutionary parallelism supports the idea that low to intermediate levels of pleiotropy are more favorable for adaptation [9, 73].

In a concluding remark, we would like to emphasize that although pleiotropy and ancestral variation were found to be significantly associated with the degree of parallelism of gene expression evolution, it is likely that additional factors also contribute. For example, diminishing return epistasis, population size (drift), and the contribution of gene expression to the selected traits may also affect the parallelism of gene evolution. Further investigation of other potential factors will continue to advance our understanding of parallelism and the predictability of evolution.

## Acknowledgments

We thank Viola Nolte for preparing all RNA-Seq libraries and supervising the maintenance of the evolution experiment. We thank Dagný Ásta Rúnarsdóttir for the suggestions on the early version of the manuscript and all member of the Institut für Populationsgenetik for discussion. Ana Marija Jakšić, Neda Barghi, François Mallard and Kathrin Otte performed the common garden experiment. Illumina sequencing was done at the VBCF NGS Unit (www.vbcf.ac.at). This research was funded in whole or in part by the Austrian Science Fund (FWF) [W1225, P32935] and the European Research Council (ERC) [ArchAdapt]. For open access PURPOSES, the author has applied a CC BY public copyright license to any author-accepted manuscript version arising from this submission..

## Author contribution

W.Y.L, S.K.H. and C.S. conceived the study. A.F. contributed statistical advice, in particular for the causal analysis. W.Y.L. performed the data analysis. S.K.H. performed the simulations.

W.Y.L. and C.S. wrote the manuscript. W.Y.L., S.K.H., A.F. and C.S. revised and edited the final manuscript.

## Data accessibility statement

All sequencing data are available in European Nucleotide Archive (ENA) under the accession number PRJEB35504, PRJEB35506 and PRJEB37011. Other raw data and scripts are available on github (https://github.com/cloudweather34/pleiotropy).

