## Supplementary information for "Pleiotropy increases parallel selection signatures during adaptation from standing genetic variation"

### Supplementary text

Data transformation and standardization

As described in the main text, we quantified the evolutionary parallelism with 1/F and the estimates were log transformed to fit normal assumption.

$$Pa = \log(1/F) \tag{1}$$

We measured the ancestral variation of gene expression with log-transformed squared biological coefficient of variation based on the individual expression data of each gene for the ancestral population.

$$A = \log(BCV^2) \tag{2}$$

The pleiotropy of a gene is approximated with the tissue-specificity of its expression and network connectivity. For each measure, 10 bins of equal sizes were derived based on deciles and genes in different bins are considered to exhibit different extent of pleiotropy.

$$Pl = j, \quad when\ \widehat{Pl}_{10(j-1)\%} < \widehat{Pl}_i \leq \widehat{Pl}_{10(j)\%} \tag{3}$$

Where $j$ = 1, 2, 3…, 10; $\widehat{Pl}_i$ is 1-τ or connectivity of a gene; i = 1, 2, 3…, n; n is the total number of genes.

We standardized all three transformed variables (Parallelism (Pa), Pleiotropy (Pl), Ancestral variation (A)) by subtracting the mean and dividing by the standard deviation for the causal and regression analysis.

Exact forms of likelihood for causal model

To uncover the causal relationship between the genetic variables of interest (i.e., evolutionary parallelism (Pa), ancestral variation (A) and pleiotropy (Pl)), we represent the correlation structures in five different models considered in the main text (Figure 4) with Gaussian likelihoods and use Bayesian Information Criteria (BIC) to determine which model is best supported by the data. Similar approach has been taken previously to infer causality among genetic features of interest [1].

The parameterization and likelihoods for each model over all putatively adaptive genes are given by:

$$\mathcal{L}(Model\,I) = \prod_{i=1}^{n}\sum_{j=1}^{k} P(Pl_j)\,l(\theta; A_i|Pl_j)\,l(\theta; Pa_i|A_i)$$

$$= \prod_{i=1}^{n}\sum_{j=1}^{k} P(Pl_j)\frac{1}{\sqrt{2\pi\sigma_A^2}}\exp\left(-\frac{\left(A_i-\mu_{A_{Pl_j}}\right)^2}{2\sigma_A^2}\right)\frac{1}{\sqrt{2\pi\sigma_A^2(1-\rho^2)}}\exp\left(-\frac{\left(Pa_i-\mu_{Pa}-\rho\frac{\sigma_A}{\sigma_{Pa}}(A_i-\mu_A)\right)^2}{2\sigma_{Pa}^2(1-\rho^2)}\right) \quad (4)$$

$$\mathcal{L}(Model\,II) = \prod_{i=1}^{n}\sum_{j=1}^{k} P(Pl_j)\,l(\theta; A_i|Pl_j)\,l(\theta; Pa_i|Pl_j)$$

$$= \prod_{i=1}^{n}\sum_{j=1}^{k} P(Pl_j)\frac{1}{\sqrt{2\pi\sigma_A^2}}\exp\left(-\frac{\left(A_i-\mu_{A_{Pl_j}}\right)^2}{2\sigma_A^2}\right)\frac{1}{\sqrt{2\pi\sigma_{Pa}^2}}\exp\left(-\frac{\left(A_i-\mu_{Pa_{Pl_j}}\right)^2}{2\sigma_{Pa}^2}\right) \quad (5)$$

$$\mathcal{L}(Model\,III) = \prod_{i=1}^{n}\sum_{j=1}^{k} P(Pl_j)\,l(\theta; A_i|Pl_j)\,l(\theta; Pa_i|A_i, Pl_j)$$

$$= \prod_{i=1}^{n}\sum_{j=1}^{k} P(Pl_j)\frac{1}{\sqrt{2\pi\sigma_A^2}}\exp\left(-\frac{\left(A_i-\mu_{A_{Pl_j}}\right)^2}{2\sigma_A^2}\right)\frac{1}{\sqrt{2\pi\sigma_{Pa}^2(1-\rho^2)}}\exp\left(-\frac{\left(Pa_i-\mu_{Pa_{Pl_j}}-\rho\frac{\sigma_A}{\sigma_{Pa}}(A_i-\mu_A)\right)^2}{2\sigma_A^2(1-\rho^2)}\right) \quad (6)$$

$$\mathcal{L}(Model\,IV) = \prod_{i=1}^{n}\sum_{j=1}^{k} P(Pl_j)\,l(\theta;\ A_i)\,l(\theta; Pa_i|A_i, Pl_j)$$

$$= \prod_{i=1}^{n}\sum_{j=1}^{k} P(Pl_j)\frac{1}{\sqrt{2\pi\sigma_A^2}}\exp\left(-\frac{(A_i-\mu_A)^2}{2\sigma_A^2}\right)\frac{1}{\sqrt{2\pi\sigma_{Pa}^2(1-\rho^2)}}\exp\left(-\frac{\left(Pa_i-\mu_{Pa_{Pl_j}}-\rho\frac{\sigma_A}{\sigma_{Pa}}(A_i-\mu_A)\right)^2}{2\sigma_A^2(1-\rho^2)}\right) \quad (7)$$

$$\mathcal{L}(Model\,V) = \prod_{i=1}^{n}\sum_{j=1}^{k} P(Pl_j)\,l(\theta;\ A_i)\,l(\theta; Pa_i)$$

$$= \prod_{i=1}^{n}\sum_{j=1}^{k} P(Pl_j)\frac{1}{\sqrt{2\pi\sigma_A^2}}\exp\left(-\frac{(A_i-\mu_A)^2}{2\sigma_A^2}\right)\frac{1}{\sqrt{2\pi\sigma_{Pa}^2}}\exp\left(-\frac{(Pa_i-\mu_{Pa})^2}{2\sigma_{Pa}^2}\right) \quad (8)$$

Where i indicates each gene, j represents different pleiotropic level, n is the total number of putatively selected genes and k equals to 10 in this study. For each likelihood model, the corresponding likelihood is maximized, and parameters are estimated using standard maximum likelihood methods.

The BICs are then computed for each model as following:

$$\mathrm{BIC} = -2\ln(\mathcal{L}_i) + k_i ln(n) \tag{9}$$

Where $\mathcal{L}_i$ are the likelihood for model I-III, $k_i$ is the corresponding number of free parameters and n is total number of putatively selected genes.

The model with the smallest BIC value is identified as the best supported model.

Regression models including the effects of two measures of pleiotropy on ancestral variation/parallelism

To examine whether two measures of pleiotropy explain ancestral variation/parallelism through common or independent features, we performed two additional regression analysis (eq. 10 and 11). All variables were transformed and standardized (see supplementary information).

First, to understand whether two pleiotropy measures explain the parallelism directly through common or independent features, we fit the regression model across all putatively adaptive genes as following:

$$y_{Pa} = \beta_A x_A + \beta_{TS} x_{TS} + \beta_{NC} x_{NC} + \varepsilon \tag{10}$$

Where $y_{Pa}$ stands for parallelism, $x_A$ is ancestral variation, $x_{TS}$ is 1-$\tau$, $x_{NC}$ is network connectivity and $\varepsilon$ is random error. $\beta_A$ , $\beta_{TS}$ and $\beta_{NC}$ are the regression coefficients corresponding to $x_A$ , $x_{TS}$ and $x_{NC}$ , respectively. The regression coefficients for each corresponding variable were shown in Table S1.

Second, to understand whether two pleiotropy measures explain ancestral variation through the common or independent feature, we fit another regression model across all putatively adaptive genes as following:

$$y_A = \beta_{TS} x_{TS} + \beta_{NC} x_{NC} + \varepsilon \tag{11}$$

Where $y_A$ stands for ancestral variation, $x_{TS}$ is 1-$\tau$, $x_{NC}$ is network connectivity and $\varepsilon$ is random error. $\beta_{TS}$ and $\beta_{NC}$ are the regression coefficients corresponding to $x_{TS}$ and $x_{NC}$, respectively. The regression coefficients for each corresponding variable were shown in Table S2.

**Supplementary figure**

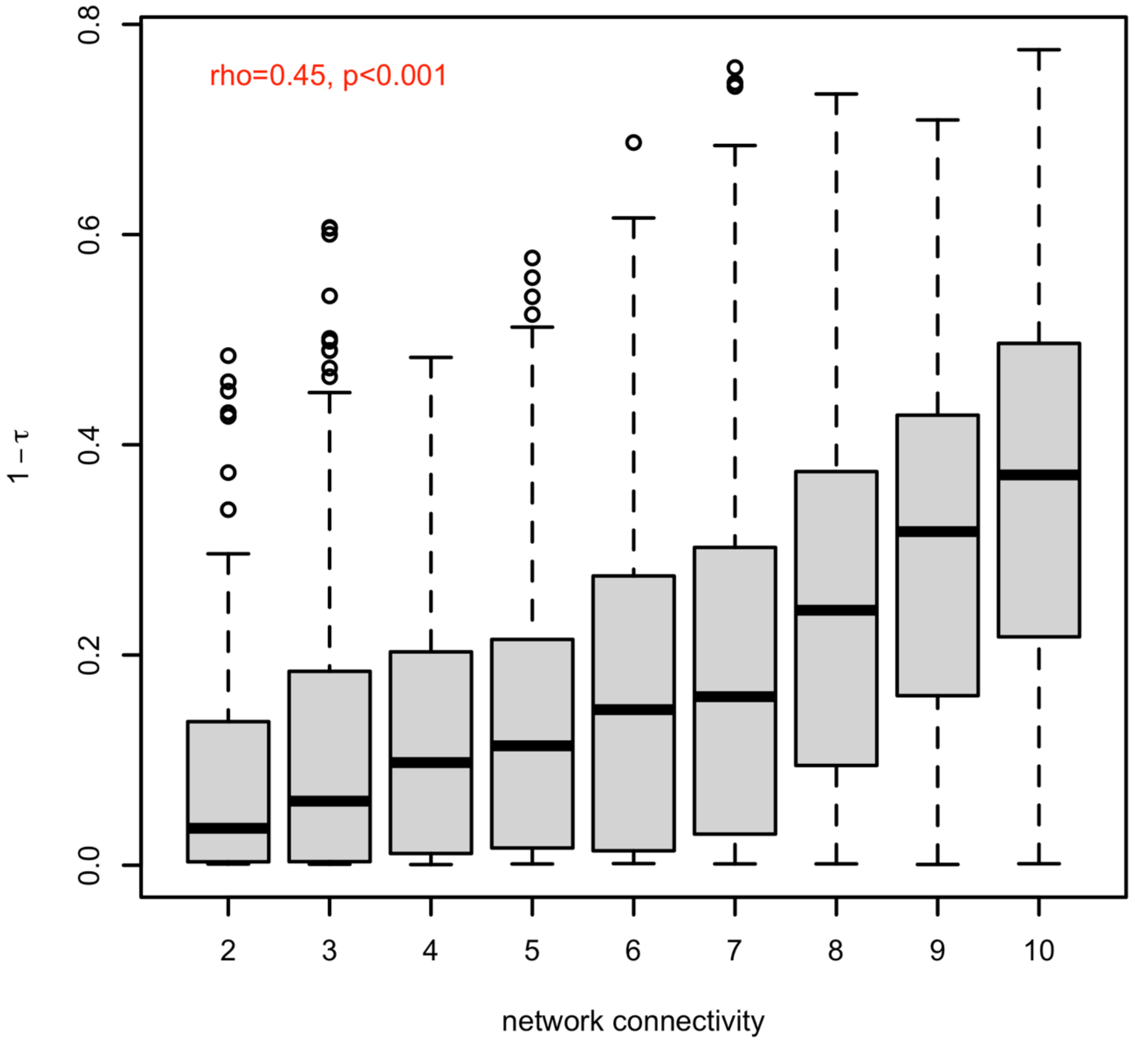


**Supplementary Figure 1. Positive correlation between two measures of pleiotropy.** The substantial correlation (rho=0.45, p.value<2.2e-16) between network connectivity (x-axis) and 1- $\tau$ (y-axis) suggest that both estimates are capturing similar information representing the pleiotropy.

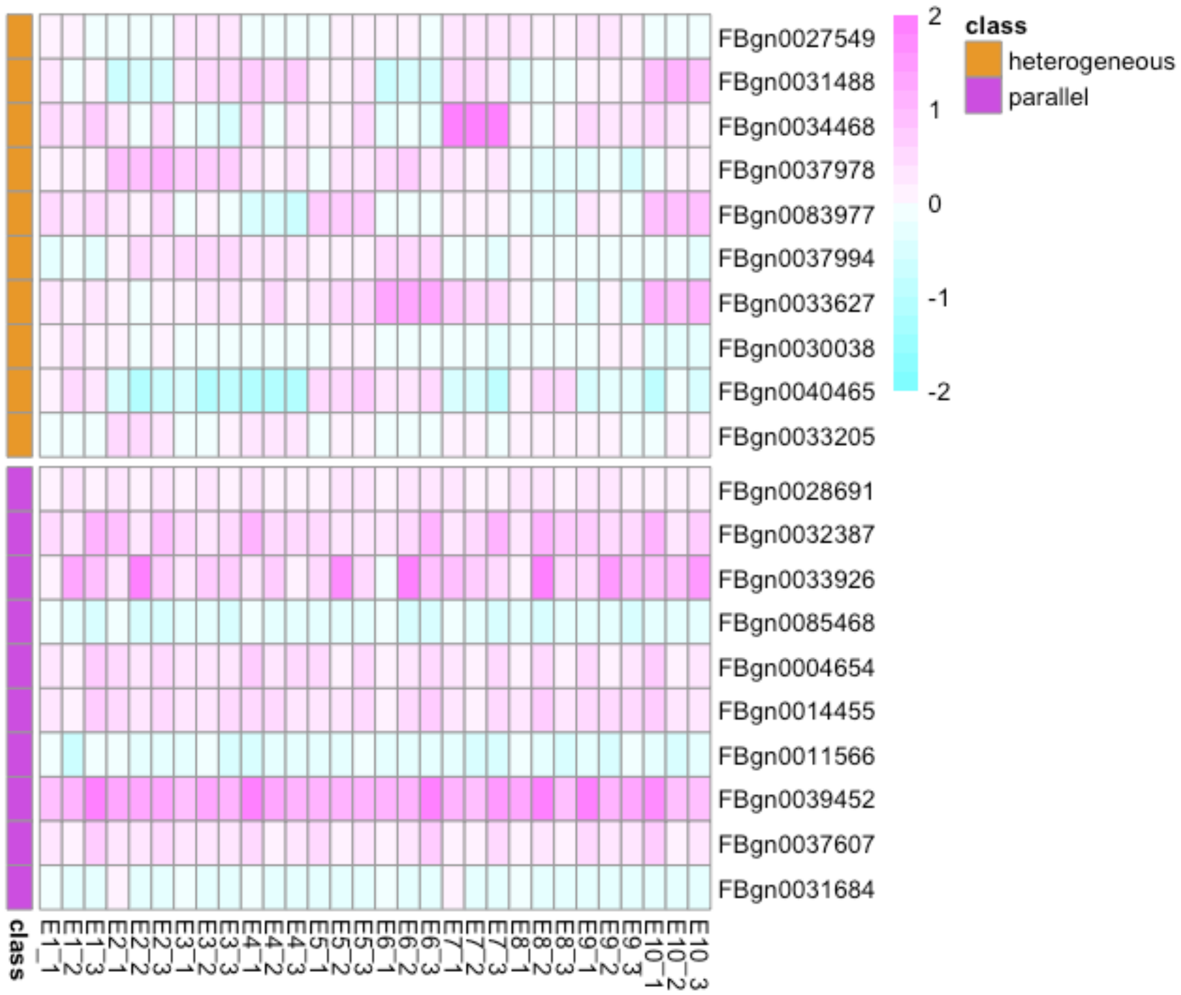


**Supplementary Figure 2. Evolutionary changes of gene expression across 10 evolved populations.** The figure compares the $\log_2$FC of top 10 genes with the most parallel evolutionary responses (genes with the top 10 highest 1/F values, shown in pink) and the top 10 genes with the most heterogeneous evolutionary responses (genes with the lowest 1/F values, shown in orange). As illustrated in the figure, genes with higher 1/F values display more parallel evolutionary responses ($\log_2$FC) across the 10 evolved populations, while genes with lower 1/F values exhibit more variation in their evolutionary responses.

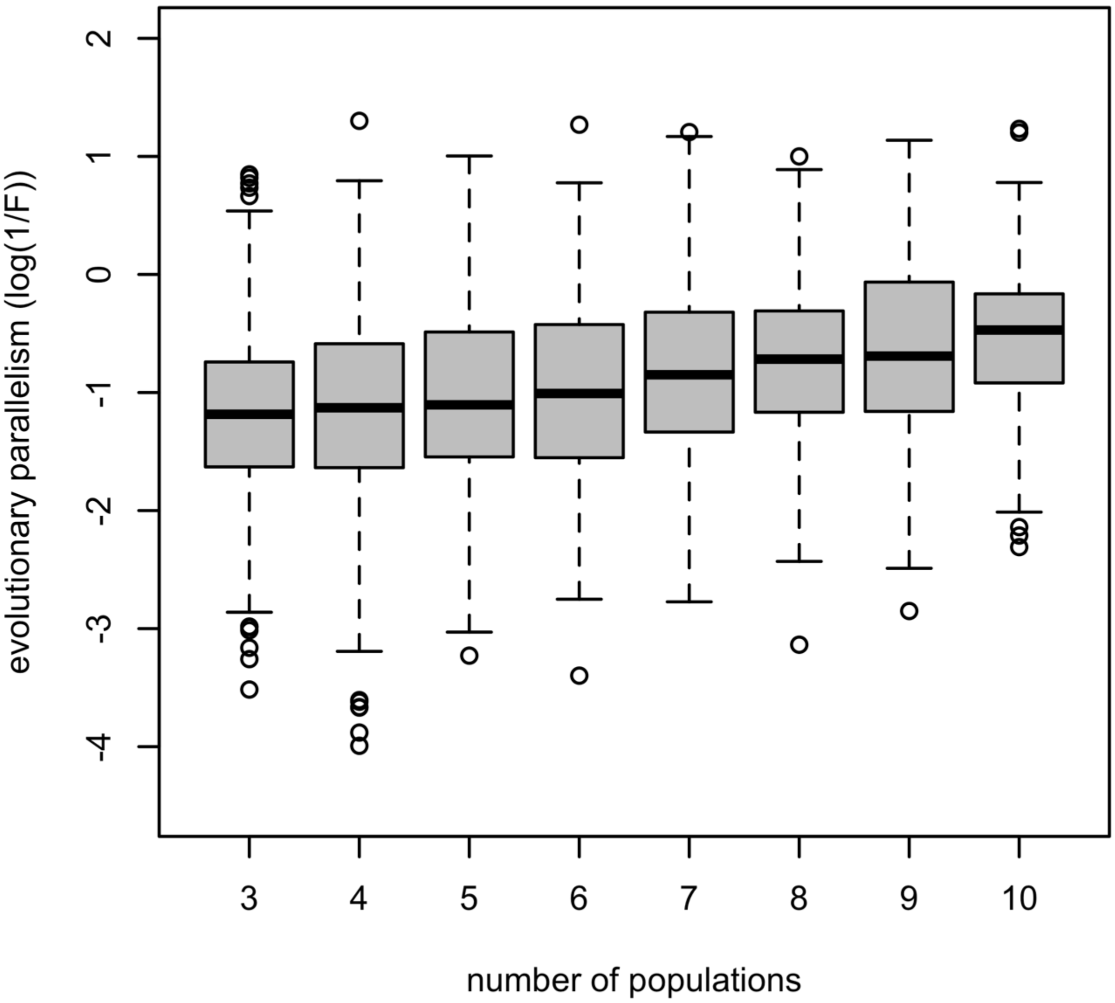


**Supplementary Figure 3. Association between parallelism of a gene (log(1/F); y-axis) and the number of evolved populations in which it is significantly differentiated from the ancestral population (x-axis).** A significant positive correlation between them (rho=0.22, p.value<5.7e-10) suggests that genes being detected in more evolved populations have higher expression parallelism and vice versa.

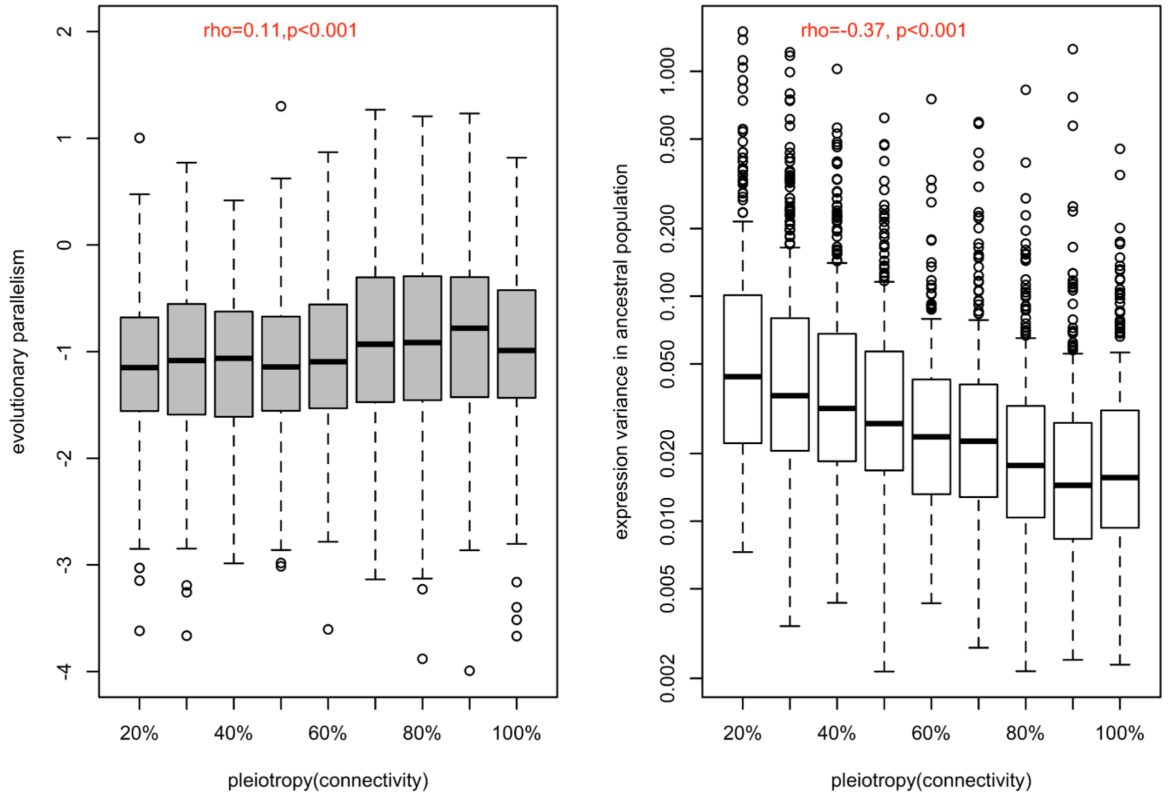


**Supplementary Figure 4. Correlation between the strength of pleiotropy (network connectivity) with the evolutionary parallelism (a) and with the ancestral variation (b). a.** The distribution of evolutionary parallelism of different genes is shown in boxplots binned by their strength of pleiotropy (network connectivity). The strength of pleiotropy was positively correlated with evolutionary parallelism (rho=0.11, p.value<5.6e-07). **b.** The distribution of expression variance in ancestral population of different genes is shown in boxplots binned by their strength of pleiotropy. The strength of pleiotropy is negatively correlated with the ancestral variation in gene expression (rho=-0.37, p.value<2.2e-16). Both results were consistent with the finding of the other measure of pleiotropy, tissue specificity.

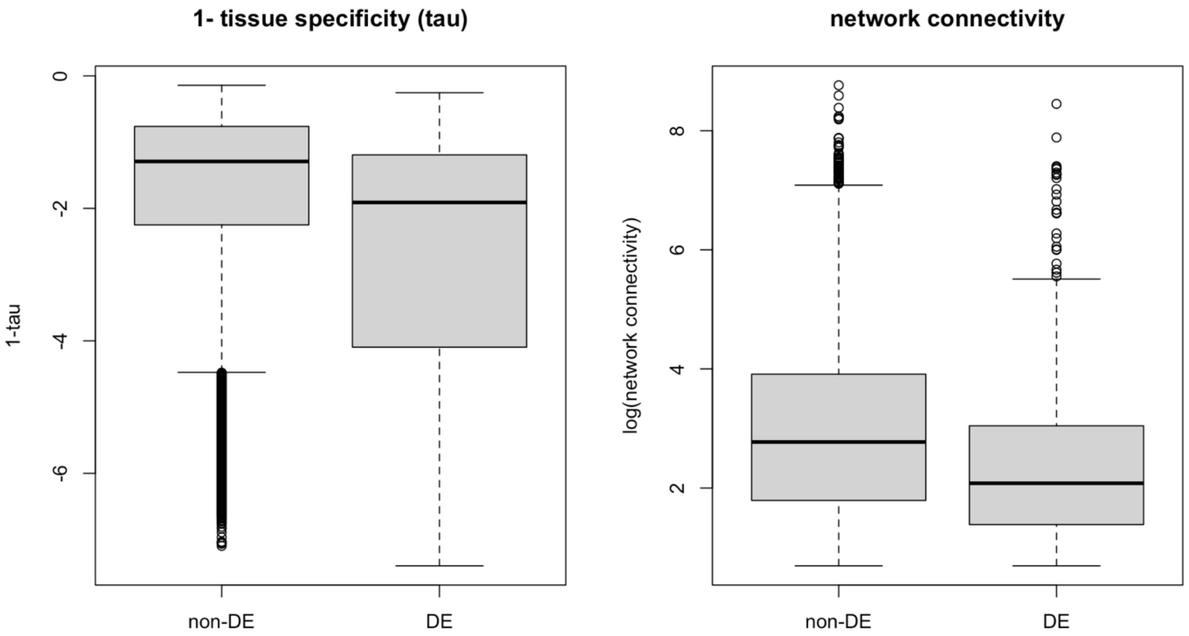


**Supplementary Figure 5. Strength of pleiotropy between DE and non-DE gene groups using (a) 1-tissue specificity and (b) network connectivity.** Genes with no evolutionary response (non-DE) have larger pleiotropic effects than genes with evolutionary responses (DE), defined here as genes showing significant expression changes in the same direction between ancestral and evolved populations in at least three populations. The same pattern is seen for both measures of pleiotropy, tissue specificity and network connectivity, suggesting that extremely high levels of pleiotropy may constrain evolutionary responses.

**Table S1. Regression of evolutionary parallelism (response) on both measures of pleiotropy (1-$\tau$ and network connectivity) and ancestral variation. Coefficient estimates and hypothesis test ($\beta = 0$) results are provided.**

| Variable | $\beta$ | F-statistics | p-value |
|---|---|---|---|
| **Ancestral variation** | -0.16 | 76.7227 | <2.2e-16*** |
| **1-$\tau$** | 0.13 | 26.4172 | 3.055e-07*** |
| **Network connectivity** | -0.03 | 1.1288 | 0.2882 |

**Table S2. Regression of ancestral variation (response) on both measures of pleiotropy (1-$\tau$ and network connectivity). Coefficient estimates and hypothesis test ($\beta = 0$) results are provided.**

| Variable | $\beta$ | F-statistics | p-value |
|---|---|---|---|
| **1-$\tau$** | -0.236 | 258.581 | <2.2e-16*** |
| **Network connectivity** | -0.239 | 95.079 | <2.2e-16*** |
